# Screening analysis of light-responsive phenotype of leaf bacteria

**DOI:** 10.1101/2025.04.16.649212

**Authors:** Hiroyuki Iguchi, Michihiro Kanagasaki, Saya Ishii, Kazuma Taguchi

**Affiliations:** Department of Agriculture and Food Technology, Faculty of Bioenvironmental Sciences, Kyoto University of Advanced Science

**Keywords:** light response, pigment, stress resistance, epiphyte, plant colonization

## Abstract

The phyllosphere is a harsh environment in which microorganisms experience various stresses, many of which are attributed to sunlight. To understand the adaptation strategy of bacteria to the phyllospheric environment under sunlight, we examined light-responsive phenotypes by cultivating leaf bacterial isolates under light. Seven bacterial strains belonging to the genera *Massilia, Kocuria, Gordonia, Agromyces, Microbacterium*, and *Arthrobacter* were identified to enhance the production of yellow or red carotenoid pigments under light exposure. Light-responsive phenotypes were further analyzed in *Massilia* sp. ST22. This bacterium was confirmed to have the ability to colonize as well as produce pigments on plants. Analysis of several forms of stress resistance revealed that light positively affected UV resistance and negatively affected resistance to osmotic pressure and oxidative stress. These results uncover the light-responsive phenotypes of the leaf bacteria and indicate that light regulates several bacterial abilities necessary for plant colonization.

## Introduction

In natural environments, microorganisms encounter various stresses, and their successful growth or survival requires adaptation to these stresses. In particular, the aerial part of plants (phyllosphere) is considered a hostile environment for microbial colonization (1, 2). The leaf surface conditions are intensely affected by climate changes (i.e., sun, rainfall, and wind), which can alter temperature, humidity, osmotic pressure, and light (UV) intensity. Furthermore, plants release antimicrobial compounds and limited nutrients. Despite the presence of these severe stresses, many microorganisms including bacteria, and fungi colonize the phyllosphere, and some play important roles in plant growth, plant health, and the carbon and nitrogen cycle (1-5). Thus, to better understand microbial ecology or implement phyllospheric microorganisms in plant cultivation, their ability to effectively colonize plant leaves has been studied (6-8). *Methylobacterium* species can utilize methanol, a niche carbon compound released from leaves, that is believed to allow this bacterial species to become dominant on leaves (9). Other analyses showed that leaf colonization requires flagella for localization and the production of trehalose, catalase, and pigment for protection from stresses (10-13).

A major influencer of the leaf surface environment is sunlight, which affects temperature, light energy, humidity, and plant metabolism. During daytime, the intensity of sunlight is altered irregularly by shading and regularly in a circadian manner. The leaf surface area is directly irradiated by sunlight, while some light energy and wavelengths are absorbed by the surface layer in soil and water environments (14, 15). The high temperature, desiccation, and DNA injury due to sunlight are harmful to microbial cells.

If microorganisms perceive changes in light intensity as a signal, they may effectively combat cognate stresses before the stresses reach a damaging level. Some bacteria employ light receptor proteins, which include LOV, BLUF, PYP, and cryptochrome for blue wavelength, and phytochrome for red wavelengths (16). Light and its cognate receptor were reported to regulate the functions necessary for plant colonization, i.e., motility, attachment, and exopolysaccharide production (17-20). However, these studies on light response used a limited panel of bacteria such as plant pathogens. The bacterial population on leaves varies widely, mainly consisting of *Proteobacteria, Bacteroidetes, Firmicutes*, and *Actinobacteria* (2, 21). Thus, knowledge of the light response of the entire leaf bacterial population is lacking, and whether leaf bacteria use light signals to facilitate growth on plants remains unknown.

In this study, we collected leaf bacterial isolates from natural environments and analyzed the phenotypes under light culturing conditions. We identified light-induced pigment production in leaf bacteria belonging to six genera. We further revealed that several forms of stress resistance of one bacterium, *Massilia*, altered under the light condition. The findings of the current study provide new information on the strategies used by leaf bacteria to grow under sunlight.

## Materials and Methods

### Sampling of plant leaves

Plant leaves were sampled in Kyoto, Hyogo and Osaka prefectures in Japan. The sampling period occurred from spring to summer in 2015, 2016, and 2017.

### Screening of light-responsive bacteria

Each leaf sample and 1 mL of sterilized water were placed in a polypropylene tube. The tube was shaken for 1 hour, and then a portion of the suspension was spread on an agar plate containing 0.5% (w/v) cycloheximide. We employed two media, commonly used for plant bacterial cultivation (22, 23): R2A broth (Nihon Pharmaceutical, Tokyo, Japan) was used in 2015 and 2016, and 2-fold-diluted tryptic soy broth (TSB; Becton, Dickinson and Company, NJ, USA) was used in 2017. The dilution of TSB was performed for the modulation of the bacterial growth rate. Colonies were collected from the agar plates to make a master plate, which was incubated at 30ºC. The collected colonies were streaked on a fresh plate containing the same agar media (1.5% agar) and cultivated at 30ºC under both dark and light (PPFD, 3 μmol m^−2^ s^−1^) conditions, the latter of which was generated with a fluorescent light (FL20SW lamp; Toshiba, Tokyo, Japan). After 2 days of cultivation, the two plates (dark and light conditions) were visually compared.

### 16S rRNA gene sequencing and accession numbers

The 16S rRNA gene was amplified by PCR with genomic DNA, the 27f-1492r primer set (24), and Blend Taq (Toyobo, Osaka, Japan), and then sequenced. These sequences were submitted to GenBank and assigned the following accession numbers: LC416392–LC416398.

### Spectral analysis

The agar plates on which the bacteria were streaked were cultivated in the dark and under fluorescent light (PPFD, 3 μmol m^−2^ s^−1^). To extract pigments, 0.5 mg (wet weight) of the colonies collected from the plates was mixed with 1 mL of methanol or acetone (for strain S136). The suspensions were incubated for 1 hour at room temperature or 50ºC (for strain S138), and then the supernatant generated by centrifugation was analyzed by a spectrophotometer (UV-2550; Shimadzu, Kyoto, Japan).

### HPLC analysis

Strain ST22 was grown on *Arabidopsis* plants by seed inoculation describes below, or on R2A agars. Cells of strain ST22 were harvested from the R2A liquid culture and washed and resuspended with water. *Arabidopsis thaliana* Col-0 seeds were sterilized with 70% ethanol. The seeds were incubated in 1 mL cell suspension (each OD_600_=0.1) with shaking on a rotator for 2 hours. The seeds were sown on Murashige and Skoog medium including vitamins (Duchefa, Haarlem, Netherlands) and supplemented with 0.25% sucrose and 0.8% agar. The plants enclosed in petri dishes were gnotobiotically incubated in a plant growth chamber (Nippon Medical & Chemical Instruments, Osaka, Japan) at 25°C (14 hours light/10 hours darkness) for 3 weeks. The light condition was prepared with two fluorescent lights (FHF32EX-N-HX-S lamp, NEC, Tokyo, Japan; FHF32N-EDL lamp, Toshiba), and the light intensity (PPFD) at the petri dishes was 110 μmol m^−2^ s^−1^.

The bacterial cells on the plant were collected from the aerial parts of six plants by suspension with water followed by centrifugation. The collected bacterial cells from plants or agars were mixed with methanol to extract the pigments, and the supernatant was subjected to HPLC analysis.

HPLC analysis was performed on a UFLC system LC-20 series (Shimadzu). Each sample was separated on an ODS column (column size 2 × 150 mm, Tosoh, Tokyo, Japan), and the column flow was maintained at 0.2 mL/min with a linear gradient of 20 to 100% acetonitrile for 15 minutes. The UV/Vis detector was set at 450 nm.

### ROS (reactive oxygen species) assay

ROS production was evaluated by a dihydroethidium-based assay (25). Strain ST22 cells, grown on R2A agar in the dark or under fluorescent light (PPFD, 3 or 10 μmol m^−2^ s^−1^) for 2 days, were suspended in Dulbecco’s phosphate-buffered saline (D-PBS; Nacalai Tesque, Kyoto, Japan). The supernatant, generated by undisturbed settling for a few minutes, was collected and adjusted to OD_600_ 0.5 with D-PBS.

One hundred microliters of the cell suspension was mixed with 50 μL of 15 μM dihydroethidium (final concentration 5 μM) and incubated at 30°C for 30 minutes. The cells were collected by centrifugation and suspended in D-PBS. The fluorescence was measured using a fluorescence microplate reader (Gemini EM; Molecular Devices, CA, USA) with an excitation wavelength of 485 nm and emission wavelength of 590 nm.

### Liquid culture growth

Strain ST22 cells were grown on R2A medium at 30°C and 160 rpm in the dark or under fluorescent light (PPFD, 6 μmol m^−2^ s^−1^).

### Stress resistance assay

Strain ST22 cells, grown on R2A agar in the dark or under fluorescent light (PPFD, 3 μmol m^−2^ s^−1^) for 2 days, were suspended in water. The supernatant, generated by undisturbed settling for a few minutes, was diluted to OD_600_=1 and used for the following analysis. Results of the analysis were expressed as viability, which was calculated by comparing the CFU of stress-treated cells with that of untreated cells. Colony formation for CFU evaluation was performed in the dark. (i) UV resistance. The cell suspension was further diluted and spread on eight plates of R2A agar. Five of eight plates were exposed to UV-C light (GL10 lamp, NEC; peak spectrum, 254 nm; full width at half maximum, 7 nm) for 10 seconds, and the three remaining plates were untreated. (ii) Heat resistance. The cell suspension was divided into four tubes (each 500 μL, OD_600_=0.1), incubated at 50°C for 15 minutes, and then immediately cooled at 4°C. The appropriately diluted suspensions were spread on R2A agar. (iii) NaCl resistance. The cell suspension was divided into four tubes containing 2 M NaCl (final OD_600_=0.1), incubated for 30 minutes, and then immediately diluted with water. (iv) H_2_O_2_ resistance. The cell suspension was divided into four tubes containing 0.05 % hydrogen peroxide (each 500 μL, final OD_600_=0.1), incubated for 15 minutes, and then immediately mixed with 5 μL of 0.2 M sodium thiosulfate for neutralization.

## Results

### Screening of light-responsive bacteria from plant leaves

We investigated the light-response behavior of the bacterial isolates from plant leaves. We used R2A agar, an oligotrophic medium, to cultivate bacteria from leaf samples. We isolated 382 bacterial strains, and determined whether they have light-responsive phenotypes. Visual comparisons between agar cultures grown under dark and fluorescent light conditions revealed two strains (Fig. 1 and Table 1): strain Kn16 (identified in the first year) produced yellow pigments under the light condition, and strain ST22 (identified in the second year) exhibited enhanced yellow pigment production under the light condition. Other light-responsive phenotypes such as motility, colony morphology, and growth rate were not detected under the tested condition. Sequencing of the 16S rRNA gene (ca. 1,500 bp) revealed that strains Kn16 and ST22 both belonged to the genus *Massilia*, with a sequence difference of 3.7%. These two strains were isolated from different plant species (Table 1). Then, isolation and cultivation of leaf bacteria were carried out using TSB agar, which is a nutritious medium. Among the 224 isolated strains, we identified five strains with enhanced pigment production under the light condition (Fig. 1 and Table 1). Strains S71, S138, S821, and S1029 produced yellow pigments, while strain 136 produced orange pigment. The 16S rRNA gene sequences revealed that all five strains belonged to the phylum Actinobacteria. Based on the closest relatives, the strains were classified as *Kocuria* sp. S71, *Gordonia* sp. S136, *Agromyces* sp. S138, *Microbacterium* sp. S821, and *Arthrobacter* sp. S1029.

**Table 1.** Bacterial isolates from leaves with light-induced pigmentation.

| Strain | Isolation origin | Closest relative |
| --- | --- | --- |
| Kn16 | *Salix koriyanagi* | *Massilia timonae* UR/MT95<sup>T</sup> (98%) |
| ST22 | *Toxicodendron sylvestre* | *Massilia putida* 6NM-7<sup>T</sup> (98%) |
| S71 | *Osmanthus fragrans* var. *aurantiacus* | *Kocuria palustris* TAGA27<sup>T</sup> (100%) |
| S136 | Not identified | *Gordonia hongkongensis* HKU50<sup>T</sup> (98%) |
| S138 | *Rhododendron pulchrum* cv. *Oomurasaki* | *Agromyces aureus* AR33<sup>T</sup> (99%) |
| S821 | *Cayratia japonica* | *Microbacterium ginsengisoli* Gsoil 259<sup>T</sup> (99%) |
| S1029 | *Conyza canadensis* | *Arthrobacter histidinovorans* DSM 20115<sup>T</sup> (99%) |

Closest relative is with the type strains. The sequence identity is shown in parentheses.

**Fig. 1.**
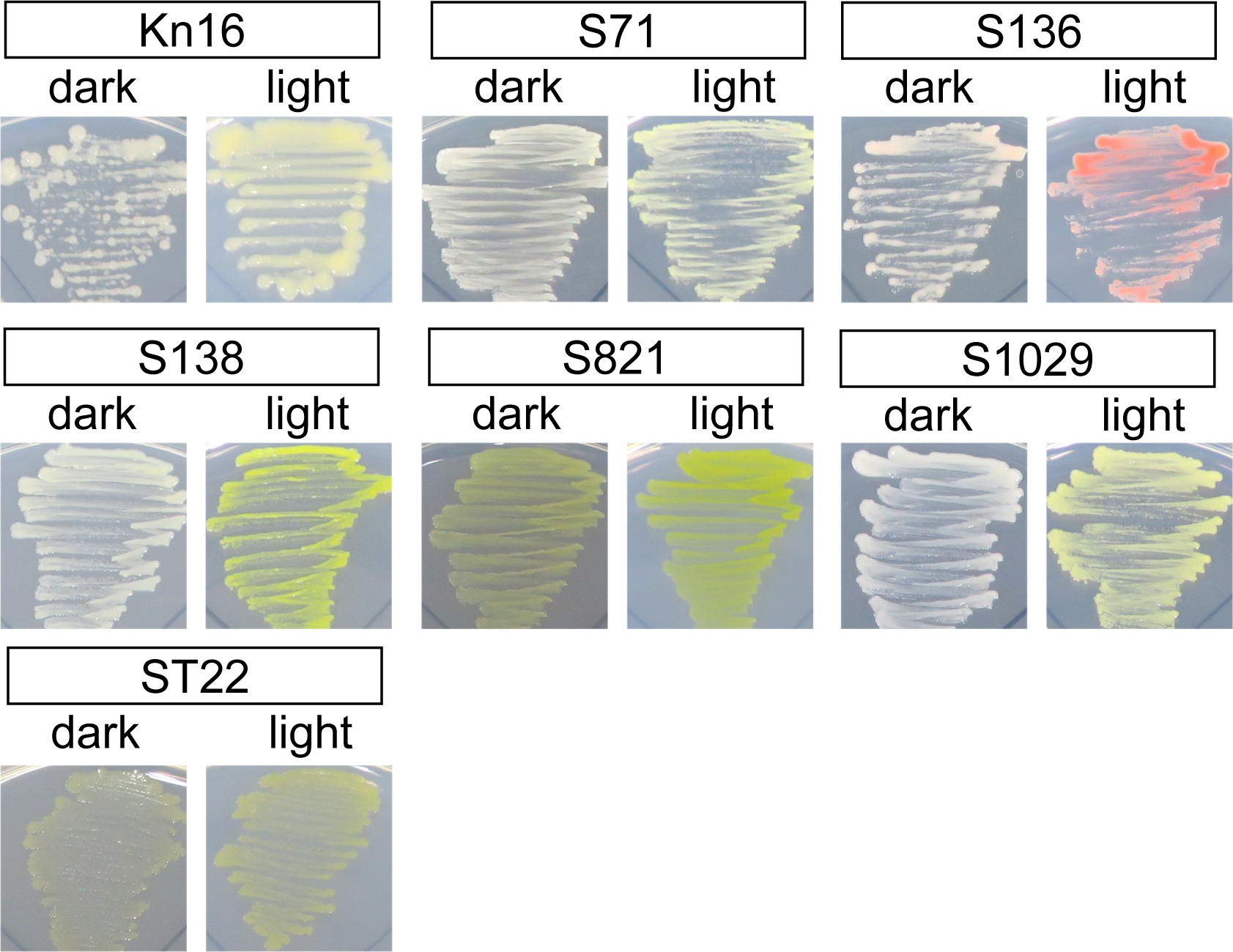
Colony of leaf bacterial isolates grown under dark or light conditions. Bacteria from leaf samples were grown on agar, and isolates were stored on a master plate. The stored bacteria were inoculated on two agar plates, and the colony phenotypes were compared between the dark and light culture conditions.

The pigment production induced under the light condition was analyzed with a spectrophotometer. The pigments were extracted from cells grown on agar under dark and light conditions. The spectra of the pigment extracts of seven strains displayed a peak representative of carotenoid, with the maximum absorption wavelength at 450 nm (Fig. 2). The pigment production was induced 2.6-to 17-fold by light exposure, based on the absorbance value at 450 nm.

**Fig. 2.**
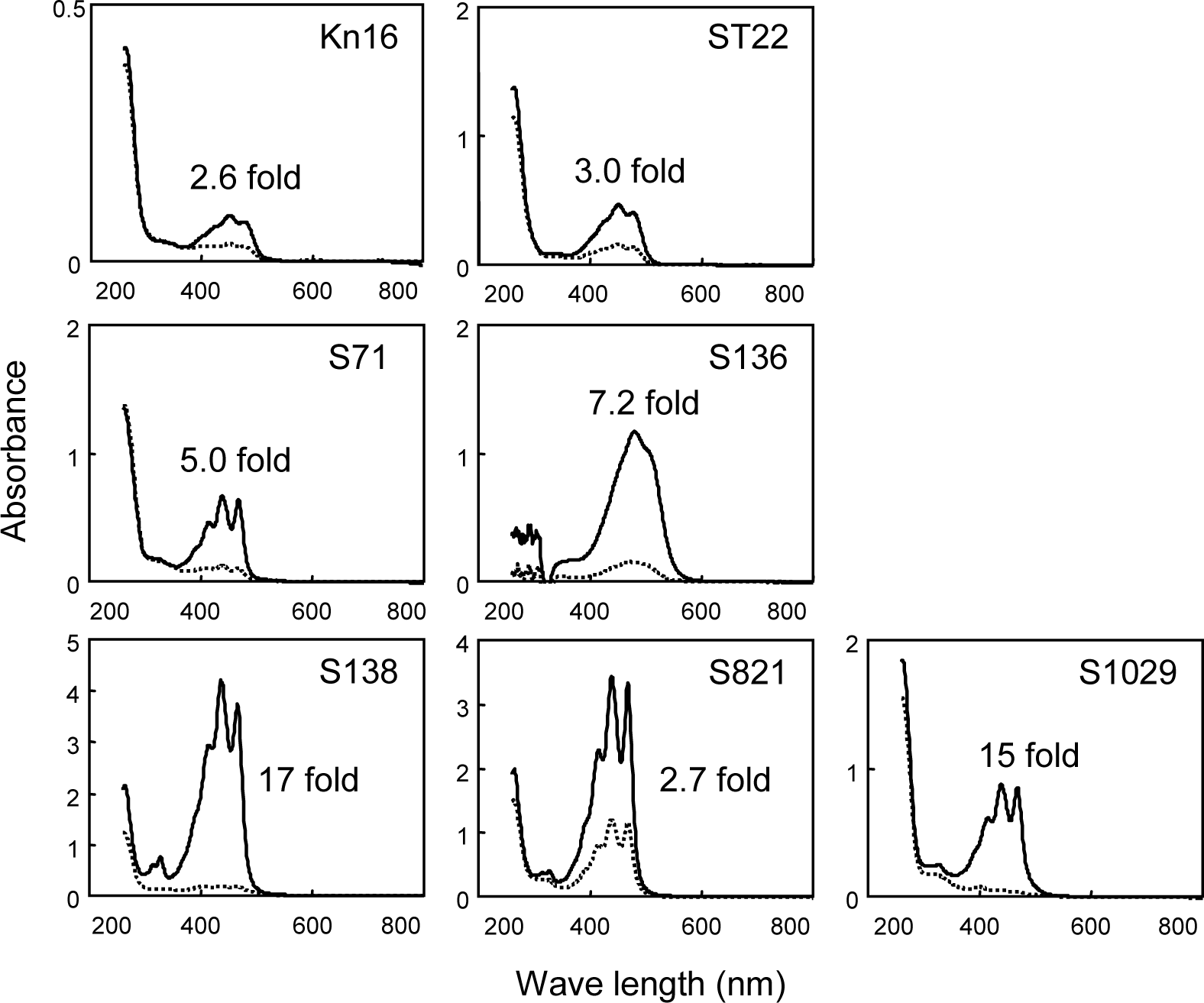
Spectrum of cell extracts of leaf bacterial isolates grown under dark (dotted line) or light (solid line) conditions. Pigments were solvent-extracted from cells grown on agar. The value indicates the fold change of the absorbance at 450 nm, which represents the maximum absorption wavelength of carotenoid pigment, between the cells grown under dark and light conditions, respectively.

### Plant colonization ability of *Massilia* sp. ST22

We determined if the pigments are produced during plant colonization. We used the isolate *Massilia* sp. ST22, as knowledge of the light-response and plant colonization of this genus is lacking. The strain ST22 was inoculated on sterilized *Arabidopsis* seeds, which were cultivated gnotobiotically under a dark/light cycle. The aqueous suspension containing the phyllospheric portions of the plants formed colonies on R2A agar, demonstrating that strain ST22 have the ability to grow on plants. Then, the pigment production on plants was analyzed by HPLC to detect the relatively small number of cells colonizing the plants. The pigments in bacterial cells collected from the plant phyllosphere were extracted with methanol. The extract of the strain showed a peak at 13.5 min (Fig. 3a), which corresponds to the pigment produced on agar (Fig. 3b), and the eluted fraction displayed a yellow color. These results showed that strain ST22 produces pigments during colonization on *Arabidopsis* plants.

**Fig. 3.**
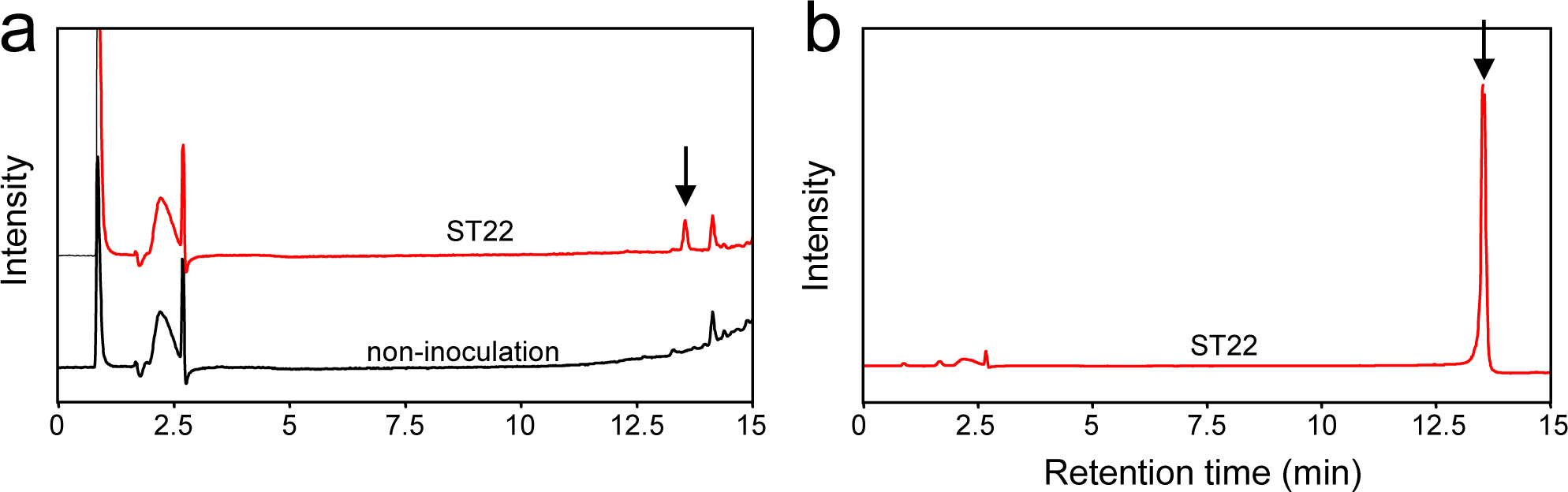
HPLC analysis of pigment production in *Massilia* sp. ST22. Pigments were extracted from bacterial cells collected from the phyllosphere of *Arabidopsis* plants (a) and agar cultures (b). An arrow indicates the peak of the pigment.

### Analysis of light-responsive, stress-resistant phenotypes of *Massilia* sp. ST22

Next, we further analyzed light-responsive phenotypes with respect to stress resistance to determine whether light regulates the plant colonization ability. First, the light conditions were set to minimize cellular damage. The ROS level was evaluated based on the dihydroethidium-based fluorescence assay for the cells grown on agar. The ROS level was comparable between light and dark conditions at a light intensity of 3 μmol m^−2^ s^−1^, whereas ROS were accumulated at a light intensity of 10 μmol m^−2^ s^−1^ (Fig 4a). This light intensity (3 μmol m^−2^ s^−1^) also induced pigment production as shown in Fig. 1 and was thus employed in the following tests with agar cultures.

**Fig. 4.**
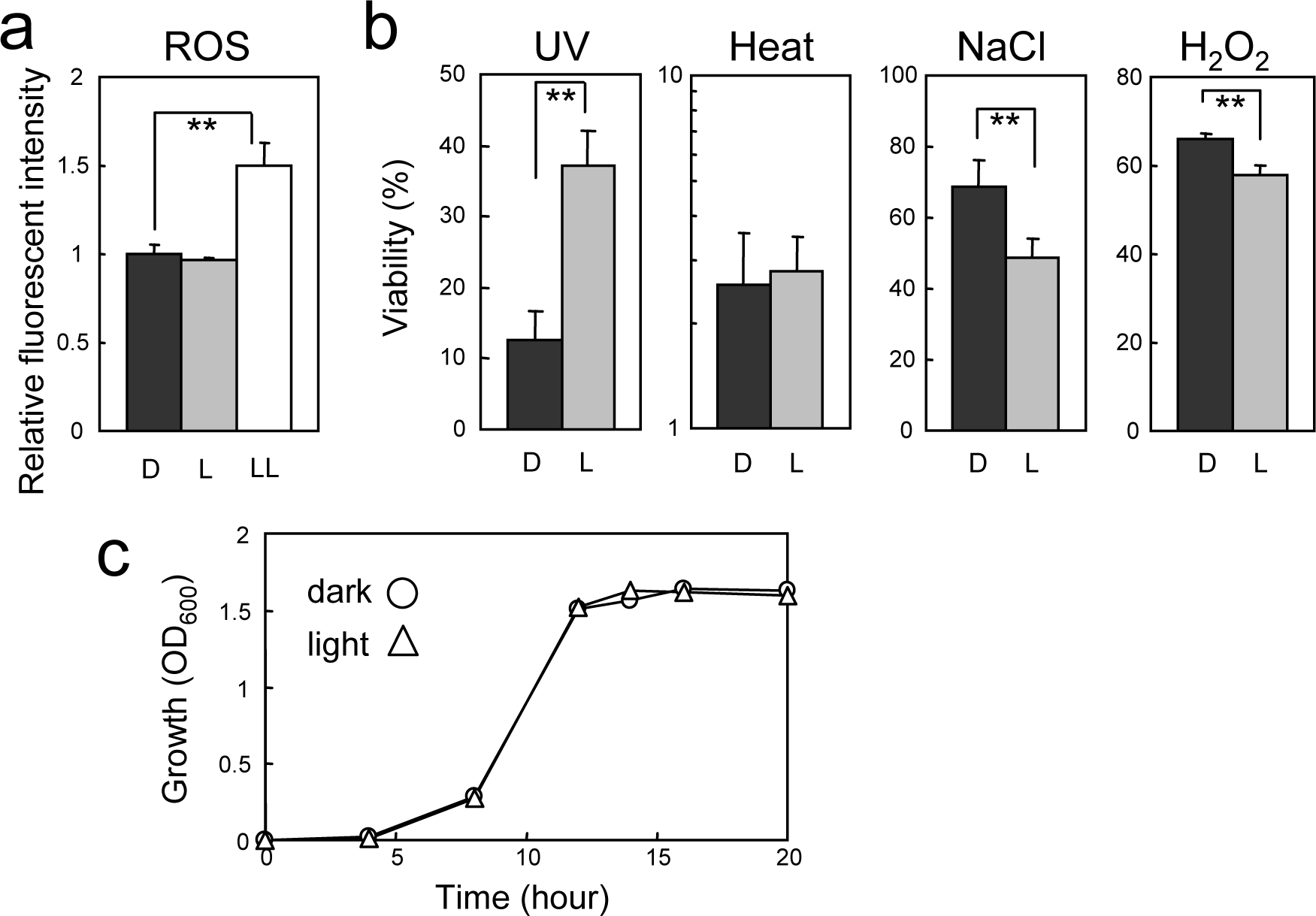
Analysis of light responsive functions in *Massilia* sp. ST22. (a) ROS generation in cells grown under dark and light conditions. Values indicate the relative fluorescent intensity based on the dihydroethidium assay. Data are provided as the means ± standard deviations (N=3). (b) Stress resistance of cells grown under dark and light conditions. The viability is expressed as the relative CFU ratio of stress-treated cells to untreated cells. Data are provided as the means ± standard deviations (N=4). D, dark condition; L, light condition at 3 mol m^−2^ s^−1^ intensity; LL, light condition at 10 mol m^−2^ s^−1^. Statistical significance was assessed using Student’s t-test. **, *P*<0.01. (c) Liquid culture growth under dark (circle) and light (triangle) conditions.

The resistance of *Massilia* sp. ST22 wild-type to several stresses was compared under dark and light conditions. It was found that UV resistance was elevated under the light condition, while the resistance to NaCl and hydrogen peroxide was decreased under the light condition (Fig. 4b). Light did not affect the heat resistance. Growth was compared in liquid culture using the flask shaker equipped with a fluorescent light (6 μmol m^−2^ s^−1^). A similar growth curve was generated for the dark and light conditions (Fig. 4c). These results indicated that light positively regulates UV resistance, and negatively regulates on resistance to osmotic pressure (NaCl) and oxidative stress (H_2_O_2_) in *Massilia* sp. ST22.

## Discussion

The phyllosphere is known to be a stressful environment, and microbial colonizers are expected to be able to effectively respond to the stresses for growth and survival. Our focus in this study was to determine whether leaf bacteria use sunlight as a signal to address the stresses present in the phyllosphere. To elucidate the light-responsive phenotypes, we cultivated a total of 606 bacterial isolates from leaves under light and dark conditions and identified seven strains that produce more pigments under light conditions (Fig. 1 and Table 1). These strains belong to the two major phyla in the phyllospheric bacterial community (Proteobacteria and Actinobacteria) (2, 21). This is the first report on the light-induced pigmentation of the two genera *Massilia* and *Kocuria*. Pigment production under light conditions has been reported in four other genera (*Gordonia, Agromyces, Microbacterium*, and *Arthrobacter*) (26-30). These six genera are widely distributed in environments such as soil, water, animals, and plants (phyllosphere and rhizosphere) (31-36). Some species of these genera were reported to not only colonize plants but also have beneficial interaction with plants (i.e., plant growth promotion) (35-40). The air pollutant biodegradation capacity of bacteria on leaf surfaces was also reported (41). Thus, these bacteria are likely to play an important ecological role in plant-associated environments.

Some isolated strains produced pigments at a low level in the dark, whereas strains S71, S138, and S1029 showed almost no pigment production in the dark, indicating strict repression of pigment synthesis (Fig. 1 and 2). Once pigments are produced intracellularly, they are usually maintained in the cells during their lifetime. Accordingly, light induced-pigmentation is likely to occur when the bacteria first encounter light. The advantages of the light-induced pigmentation system in bacteria remain unknown. One explanation is that bacteria can conserve the biological energy for pigment production in dark environments such as soils where pigment function is possibly not required. The plant pathogen *P. syringae* was reported to reversibly change its behavior for plant infection via the blue light receptor between the soil (dark) and phyllospheric (light) environments as well as day and night. Under blue light conditions, which partly represent the phyllospheric environment, motility and virulence are repressed, while attachment to plant leaves is promoted (17). At nightfall, these traits can be reversed (i.e., activation of motility and virulence) by altering gene expression levels. The defense response of plants is also regulated in a circadian manner, with higher susceptibility to infection at night (42).

The induced pigments were considered carotenoids (Fig. 2), and carotenoid pigments are known to function in protection from oxidative stresses, especially ROS generated by UV radiation (43, 44). Previous studies demonstrated that pigment-deficient bacterial mutants became more sensitive to oxidative stresses (12, 45, 46) with reduced ability to colonize plants (12, 44). The plant colonization ability encompasses many traits, including acquisition and metabolism of nutrients, response to environmental stresses (temperature, light, humidity, osmotic pressure, ROS, and antimicrobial compounds), motility, and adhesion (2). Several light-responsive phenotypes possibly relative to plant colonization were further revealed in *Massilia* sp. ST22. Light induced UV resistance (Fig. 4b), representing the close connection between light and resistance to damage caused by light. Thus, this bacterium is suspected to generate a defense system against sunlight damage by enhancing its UV resistance during daytime. Light-induced UV resistance has been observed in some microorganisms (47, 48) and probably occurs via controlling the functions of DNA repair, protein folding, and ROS scavenging at the gene expression level (43, 49-51). In the context of UV resistance, the resistance to oxidative stress, which is caused by light, was expected to be induced, but the resistance to H_2_O_2_ was decreased (Fig. 4b). The resistance to osmotic pressure (NaCl), which is unlikely to relate to light, was also found to be decreased. Thus, not all light-induced phenotypes are useful for plant colonization, and we suspect that light triggers multiple regulatory systems (i.e., positive and negative regulation).

In summary, cultivation of bacteria under light conditions has clarified the light-responsive phenotypes of seven phyllospheric bacteria. Our test conditions using agar culture clearly revealed the pigmentation phenotype. The extensive analysis of different bacterial abilities revealed that several stress-resistance phenotypes of *Massilia* sp. ST22 affected by light. This finding suggests the possibility that light-responsive phenotypes are widely distributed in phyllospheric bacteria, including those not assessed in this study. The light-responsive phenotypes such as stress resistance also should play a role in growth and survival in the phyllosphere. It is still unclear whether changes in stress resistance under the light condition occurred via sensing of light signals with photoreceptors or were triggered by light-induced cell injury. Further work is required to determine the light-responsive regulatory mechanism, including photoreceptor proteins, transcriptional factors and regulated genes. In addition, techniques controlling bacterial behavior using specific wavelengths and photoreceptors are promising research topics for future investigation (52-54). The combined knowledge of ecology and molecular biology will facilitate the application of light to control the abilities of phyllospheric bacteria, including growth property, pathogenicity, and bioactive compound production.

## Funding

This study was supported in part by a Grant-in-Aid for Young Scientists (18K14381, to H.I.), and a Grant-in-Aid for Young Scientists (B) (15K18671, to H.I.) from the Japan Society for the Promotion of Science.

## Notes

### Competing Interest Statement

The authors have declared no competing interest.

### Summary of Updates

Figure position has adjusted because the header overlapped the figures in the printed format.

